# Structurally and functionally distinct early antibody responses predict COVID-19 disease trajectory and mRNA vaccine response

**DOI:** 10.1101/2021.05.25.445649

**Authors:** Saborni Chakraborty, Joseph C. Gonzalez, Benjamin L. Sievers, Vamsee Mallajosyula, Srijoni Chakraborty, Megha Dubey, Usama Ashraf, Bowie Yik-Ling Cheng, Nimish Kathale, Kim Quyen Thi Tran, Courtney Scallan, Aanika Sinnott, Arianna Cassidy, Steven T. Chen, Terri Gelbart, Fei Gao, Yarden Golan, Xuhuai Ji, Seunghee Kim-Schulze, Mary Prahl, Stephanie L. Gaw, Sacha Gnjatic, Thomas U. Marron, Miriam Merad, Prabhu S. Arunachalam, Scott D. Boyd, Mark M. Davis, Marisa Holubar, Chaitan Khosla, Holden T. Maecker, Yvonne Maldonado, Elizabeth D. Mellins, Kari C. Nadeau, Bali Pulendran, Upinder Singh, Aruna Subramanian, Paul J. Utz, Robert Sherwood, Sheng Zhang, Prasanna Jagannathan, Gene S. Tan, Taia T. Wang

## Abstract

A damaging inflammatory response is strongly implicated in the pathogenesis of severe COVID-19 but mechanisms contributing to this response are unclear. In two prospective cohorts, early non-neutralizing, afucosylated, anti-SARS-CoV-2 IgG predicted progression from mild, to more severe COVID-19. In contrast to the antibody structures that predicted disease progression, antibodies that were elicited by mRNA SARS-CoV-2 vaccines were low in Fc afucosylation and enriched in sialylation, both modifications that reduce the inflammatory potential of IgG. To study the biology afucosylated IgG immune complexes, we developed an *in vivo* model which revealed that human IgG-FcγR interactions can regulate inflammation in the lung. Afucosylated IgG immune complexes induced inflammatory cytokine production and robust infiltration of the lung by immune cells. By contrast, vaccine elicited IgG did not promote an inflammatory lung response. Here, we show that IgG-FcγR interactions can regulate inflammation in the lung and define distinct lung activities associated with the IgG that predict severe COVID-19 and protection against SARS-CoV-2.

**One Sentence Summary:** Divergent early antibody responses predict COVID-19 disease trajectory and mRNA vaccine response and are functionally distinct *in vivo*.

---

The minority of people who develop severe COVID-19 during SARS-CoV-2 infection mount an inflammatory response that is strongly implicated in disease pathogenesis (*1–3*). The extreme inflammatory phenotype in the lungs of severe COVID-19 patients is clear from autopsy studies, but mechanisms contributing to this response are not well understood (*4–7*) IgG antibodies mediate cellular functions that are central in directing the course of disease during many viral infections. Aside from neutralizing activity, IgG antibodies that bind to virus particles or viral antigens can form immune complexes (ICs) that may have a profound impact on disease pathogenesis, especially with regard to inflammation. This is observed in some autoimmune and infectious diseases where persistent ICs drive a hyperinflammatory response that damages host tissues (*8*). A clear mechanism underlying modulation of inflammation by antibodies is via IgG interactions with activating and inhibitory Fc gamma receptors (FcγRs) on myeloid cells, which are central regulators of the inflammatory response. We and others have previously found that patients with severe COVID-19 produce a high level of afucosylated IgG antibodies that trigger inflammatory responses in primary monocytes (*9–11*). This response was dependent on Fc afucosylation, a modification that enhances affinity of monomeric IgG for the activating FcγR, CD16a, by approximately 10-fold (*12*, *13*).

Because IgG ICs can promote disease sequelae in some infections, the link between severe COVID-19 and afucosylated IgG suggests that this antibody type may have a role in the inflammatory pathogenesis of severe disease. To explore this, we first studied whether afucosylated antibody production was a consequence of, or an antecedent to, the development of more severe COVID-19. In two independent cohorts, assessed during an initial period of mild symptoms, we found that the absence of early neutralizing antibodies, together with an increased abundance of afucosylated IgG, predicted rapid progression to more severe disease. Elevated frequencies of monocytes expressing the receptor for afucosylated IgG, CD16a, also predicted more severe outcomes. To study the effect of afucosylated antibody signaling in the lungs, we developed a novel model system in which human ICs of defined composition are intratracheally administered to mice that express human FcγRs (*14*). Molecular and cellular changes that were triggered in the lung by distinct antibody signaling pathways can then be assessed by characterization of broncho alveolar lavage (BAL) fluid collected after IC administration. This model provides a physiologically relevant system to study antibody effector responses in the lung. We observed that afucosylated ICs triggered robust immune cell activation, infiltration into the lungs, and proinflammatory cytokine production that was CD16a-dependent. In contrast to natural infection, SARS-CoV-2 mRNA vaccination elicited IgG antibodies that were both highly fucosylated and sialylated. Immune complexes formed from the mRNA vaccine-elicited IgG did not trigger the cellular infiltration and the cytokines/chemokines that were associated with afucosylated IgG *in vivo*. Overall, these findings demonstrate that early production of non-neutralizing, afucosylated IgG1 was predictive of COVID-19 symptom progression; these antibodies were structurally and functionally distinct from IgG1 elicited by mRNA SARS-CoV-2 vaccination.

## Results

### Study cohorts

To study the early antibody features that correlated with different disease outcomes in COVID-19, we characterized IgG from two longitudinal cohorts of COVID-19 outpatients from Stanford Hospital Center (n=109 Cohort 1 at enrollment; n=69 Cohort 2). While these samples were collected from interventional clinical studies, we present data only from the placebo arm of both studies; thus, our findings are not impacted by the experimental treatments trialed in either study. Participants in both studies were enrolled early in infection, within three days of a positive SARS-CoV-2 PCR test. All subjects presented with mild COVID-19 and had mild symptoms at the time of enrollment, as determined by a physicians assessment (*15*). While uncomplicated resolution of mild disease occurred in the majority of subjects, a subset of patients in each cohort (n=8 in Cohort 1; n=7 in Cohort 2) developed worsening symptoms in the hours or days following enrollment. These subjects were evaluated in the emergency department and some required hospitalization; one subject succumbed to disease. We term these patients with distinct disease trajectories as “progressors” (Tables S1, S2, S3) or “non-progressors”. Progressors and non-progressors from Cohort 1 did not significantly differ by the parameters of age, weight, or sex. Progressors from Cohort 2 also did not differ based on weight or sex but were older compared to non-progressors (Table S1).

### Low early neutralizing IgG levels in progressors

The availability of samples from the date of enrollment in both studies (here termed “day 0”), when all subjects had mild disease, enabled our analysis of early antibody responses that correlated with distinct disease trajectories. We first defined the evolution of the neutralizing antibody response following SARS-CoV-2 infections using a pseudotyped vesicular stomatitis virus neutralization assay. The fifty percent pseudoviral neutralizing antibody titers (pNT_50_) were calculated for day 0, day 5, day 28, month 7, and month 10 for all subjects in the placebo arm of Cohort 1 from whom samples were available. Samples from study participants who received a SARS-CoV-2 vaccine within the study period were not evaluated. In most subjects, levels of neutralizing antibodies showed a significant increase over time, peaking by day 28. Once initiated, the antibody response was durable and persisted in most people until 7 months post-enrollment, after which there was a general decrease in neutralization by month 10 (Fig. 1A, S1A). This analysis of Cohort 1 revealed that although there was considerable heterogeneity in early neutralizing responses, those participants who would progress to more severe disease had uniformly very low or no detectible neutralizing antibodies at the study day 0/enrollment time point (Fig. 1B). Cohort 2 showed somewhat less heterogeneity in early neutralizing responses, but as with Cohort 1, neutralizing antibodies were not detected on day 0 in any of the progressors (Fig. 1B). These data were broadly consistent with studies showing a correlation between early neutralizing antibody responses and outcomes in COVID-19 (*16–19*).

**Figure 1.**
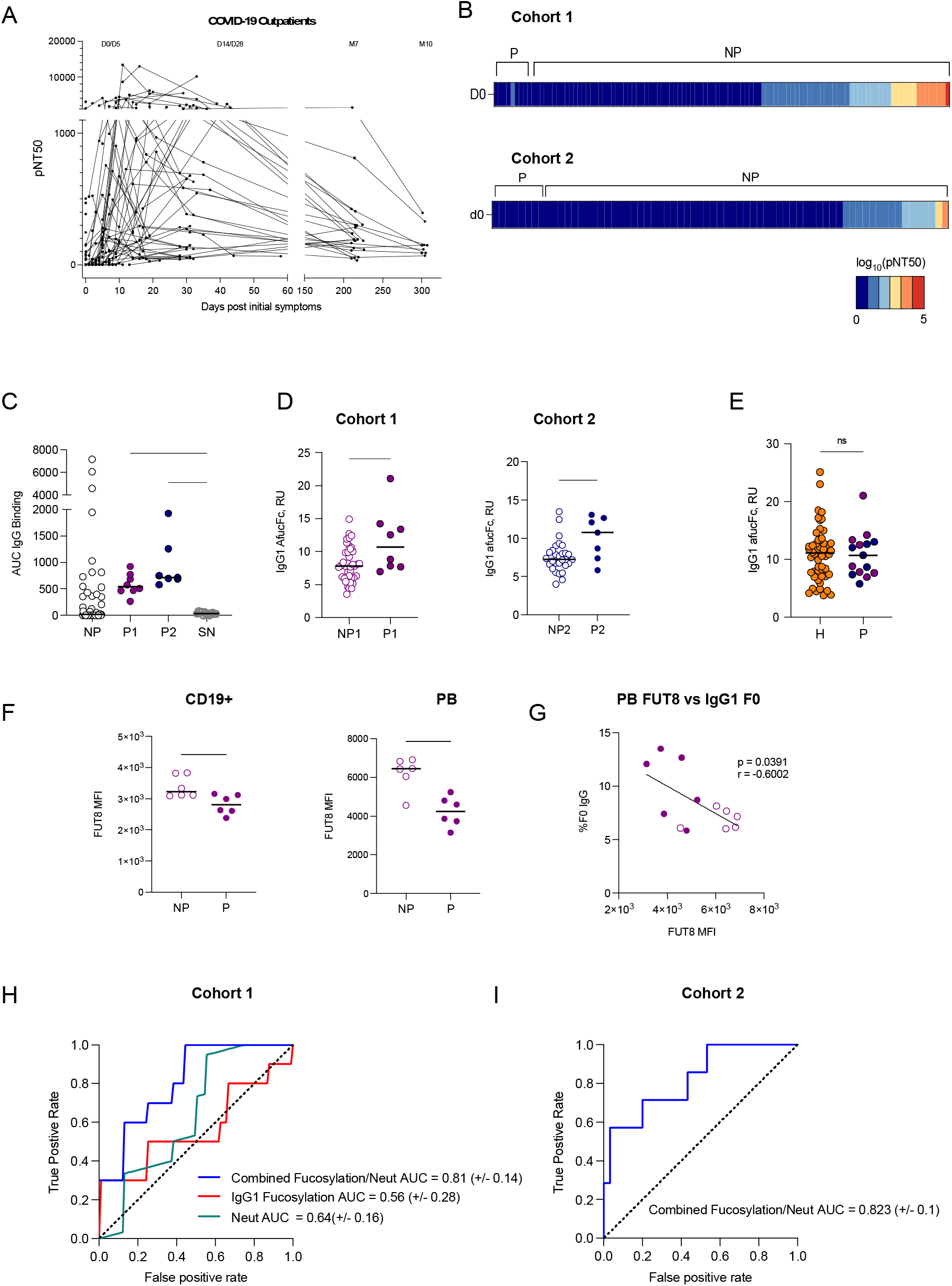
Low early neutralizing titers and elevated Fc afucosylation predict disease progression. **(A)** The kinetics of neutralizing antibody response over time in Cohort 1. Half-maximal SARS-CoV-2 pseudovirus neutralizing titers (pNT_50_) at each study time point, graphed based on days of symptoms for each participant. Samples were collected at study day 0 (D0 enrollment; n=101), 5 (D5; n=50), 14 (D14; n=33), 28 (D28; n=43), month 7 (M7; n=24) and month 10 (M10; n=9). **(B)** Heatmap of pNT_50_ data in progressors (P) (Cohort 1 n=8, Cohort 2 n=7) and non-progressors (NP) at enrollment timepoint (D0). **(C)** SARS-CoV-2 spike-binding IgG (AUC) of Cohort 1 progressors (P1, solid purple), Cohort 2 progressors (P2, solid blue), random subset of non-progressors, and historic seronegative (SN) sera. **(D)** IgG1 Fc afucosylation levels of progressors and non-progressors at enrollment timepoint (D0) of Cohort 1 (purple; progressors=P1, non-progressors=NP1), Cohort 2 progressors (blue; progressors=P2, non-progressors=NP2). **(E)** IgG1 Fc afucosylation levels in COVID-19 patients who were hospitalized (H, orange) (n=52) and combined outpatient progressors (P, Cohort 1 progressors: purple), Cohort 2 progressors: blue) (n=15). **(F)** α-1,6-Fucosyltransferase 8 (FUT8) median fluorescence intensity in total CD19+ B cells and in plasmablasts (PB) from progressors (n=6) relative to sex-matched non-progressors (n=6). **(G)** Correlation in matched samples: plasmablast expression of FUT8 and the abundance of IgG1 afucosylation. Solid and open circles represent data points from progressors and non-progressors, respectively. **(H)** Mean receiver operating characteristic (ROC) response and the area under the curve (AUC) with its standard deviation obtained using a logistic regression model using neutralization titers and IgG1 afucosylation levels. **(I)** Receiver operating characteristic (ROC) response and the area under the curve (AUC) with standard deviation obtained by testing logistic regression model on an independent Cohort 2. Median values are depicted in C-F with a solid black line. P values in (C) were calculated using Brown-Forsythe and Welch ANOVA test with Dunnett T3 correction, in (D-E) were calculated using Wilcoxon rank-sum test and in (F) using unpaired Student’s test. *P < 0.05; **P < 0.01; ***P < 0.001; ****P < 0.0001. Pearson’s correlation coefficient (r = −0.6002, p = 0.0391,) was computed in (G).

We initially reasoned that the absence of early neutralizing antibodies in progressors might have been due to earlier sampling of participants who were on a more severe disease trajectory. To evaluate this, we compared the number of symptomatic days prior to study enrollment in progressors and non-progressors. This revealed that there was no significant difference in the mean or median duration of symptoms prior to enrollment (Table S1). Thus, the kinetics of sampling did not explain this observation. Despite the absence of early neutralizing responses, SARS-CoV-2 spike-reactive IgG was clearly present in all progressors (Fig. 1C). While early neutralizing responses were not detected, progressors from whom longitudinal samples were available generally mounted neutralizing antibody responses by the later study timepoints (Fig. S1B).

### Elevated early production of afucosylated IgG in progressors

We next asked whether there were qualitative differences in the Fc structures of the IgG in progressors and non-progressors. As we had previously observed elevated anti-SARS-CoV-2 Fc afucosylation in hospitalized patients compared to outpatients (*9*), we sought to clarify whether these antibodies were produced in response to severe disease or whether they might precede the development of severe symptoms. To study this, we evaluated Fc glycosylation on antibodies present at study enrollment when all subjects had mild symptoms. Indeed, at study enrollment the progressors in both cohorts were already distinguished by elevated levels of afucosylated IgG1, comparable to the levels observed in a cohort of hospitalized COVID-19 patients in the Mount Sinai Health System (Fig. 1D, E, Table S1) (*20*). The abundance of afucosylated IgG1 in COVID-19 outpatients was not different across timepoints that were separated by approximately 200 days (Fig. S1C). These data show that production of afucosylated IgG preceded the onset of severe symptoms and afucosylated antibodies were maintained over time.

We next sought to investigate the basis of differences in antibody fucosylation. We hypothesized that differences in expression of the relevant glycosyltransferase, α-1,6-fucosyltransferase (FUT8), by antibody-secreting cells, might play a role. To investigate this, we assessed FUT8 protein levels in peripheral blood mononuclear cells (PBMCs) from progressors and non-progressors (Fig. S2A). At the time of this experiment, only PBMCs from Cohort 2 were available from the enrollment timepoint. Because we have previously observed a sex-based difference in antibody afucosylation (*9*), an equivalent number of sex-matched non-progressors were selected for this analysis. We observed no other correlations between demographic features and IgG afucosylation in either cohort (Fig. S1D). Consistent with the elevated production of afucosylated IgG by progressors, CD19+ B cells and plasmablasts from progressors expressed less FUT8 than cells from non-progressors upon enrollment (Fig. 1F). FUT8 expression within total PBMCs was comparable between groups, as was the distribution of B cell subsets, suggesting that FUT8 expression is regulated at the effector cell level (Fig. S2A, S2B, S2C). Of note, plasmablast expression of FUT8 correlated with IgG1 Fc afucosylation, supporting the hypothesis that IgG afucosylation is regulated at least in part by the expression level of FUT8 (Fig. 1G).

### Early non-neutralizing, afucosylated anti-spike IgG predict worsening symptoms in COVID-19 outpatients

To determine whether the combination of low/no neutralizing antibodies and elevated IgG Fc afucosylation was a predictor of worsening disease trajectory in mild COVID-19 patients, we next trained and evaluated a logistic regression model by using day 0 neutralization titers and afucosylated IgG frequency as input features from Cohort 1. Individually, both early neutralization titers and Fc afucosylation had low to modest predictive power to separate progressors and non-progressors, while combining the two features could separate progressors from non-progressors with higher predictive accuracy (Fig. 1H). Subsequently, the Cohort 1 data was used as the training set and the performance of the model was evaluated on an independent test dataset (Cohort 2). As shown, the combined features could discriminate divergent disease outcomes with area under the receiver operating characteristic (ROC) curve (AUC) of 0.81 (Fig. 1I). Thus, early production of reactive, afucosylated antibodies and poor serum neutralizing activity predicted progression from mild COVID-19 to more severe outcomes.

### The receptor for afucosylated IgG1, CD16a, is enriched in the myeloid compartment of progressors

In addition to afucosylated antibody production, a hallmark of patients with severe COVID-19 is inflammatory myeloid cell infiltration into the lung and excessive inflammatory cytokine production (*2, 21, 22*). These cells express the low affinity FcγRs CD32a (activating), CD32b (inhibitory) and, on some subsets, CD16a (activating). These low affinity FcγRs are engaged through avidity-based interactions when ICs are formed during infection. Depending on the magnitude of activating or inhibitory signal received upon engagement, an effector cell will respond with a proportional level of inflammatory activity (*9*). Considering that severe COVID-19 is often characterized by an aberrant effector cell activation state (*1, 23–25*), we next sought to define the expression of activating and inhibitory FcγRs on peripheral monocytes from progressors and non-progressors that might counterbalance or compound an enrichment of afucosylated IgG.

To study this, available PBMC samples collected at study enrollment were assessed for the frequency of CD16a-expressing monocyte subsets as well as their expression of all low affinity FcγRs (Fig. S3A). Notably, we found that progressors had significantly increased frequencies of total CD16a+ monocytes, CD16a+ CD14-non-classical monocytes, and CD16a+ CD14+ intermediate monocytes within the peripheral CD11c+ HLA-DR+ myeloid cell compartment compared to non-progressors upon study enrollment (Fig. 2A) (*26–28*). Further, quantitative expression analysis of CD16a within these immune cell subsets revealed higher levels of CD16a on cells from progressors, while other low affinity FcγRs (CD32a/b) were not differentially expressed (Fig. 2B, Fig. S3B). Taken together, early CD16a expression within the peripheral myeloid cell compartment predicted the development of more severe symptoms in COVID-19 outpatients (Fig. 2C,D).

**Figure 2.**
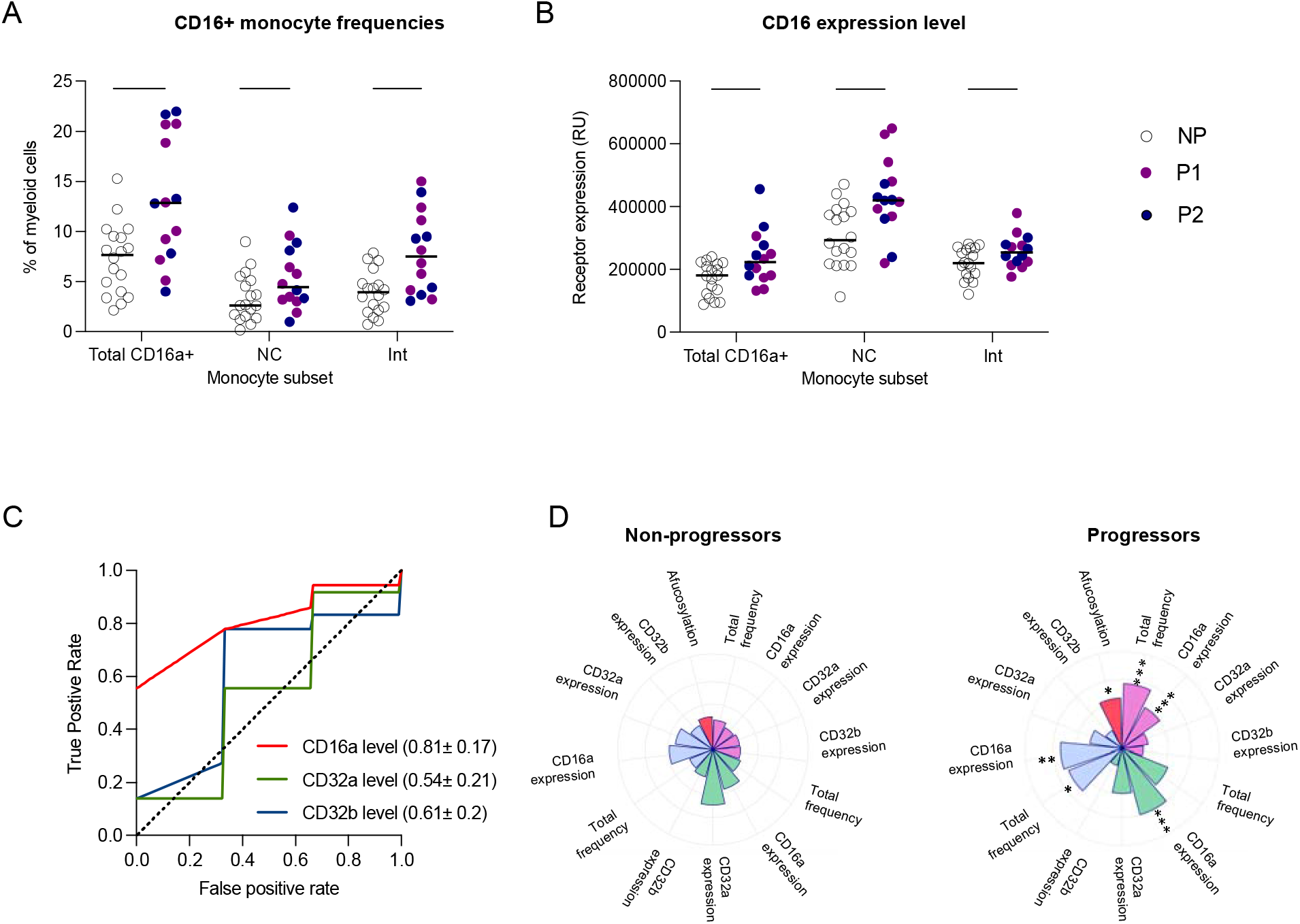
Elevated CD16a signaling potential in myeloid compartment in progressors. Enrollment time point PBMCs were characterized in progressors (n=14) and a randomly selected subset of non-progressors (n=18). Solid purple and blue circles represent data points from progressors within Cohort 1 and Cohort 2, respectively whereas open circles represent data points from non-progressors. The median values have been depicted with a black line. **(A)** Total CD16a+ monocyte, CD16a+ CD14-non-classical monocyte (NC), and CD16a+ CD14+ intermediate monocyte (Int) frequencies as percents of total CD11c+ HLA-DR+ CD3- CD19- CD56- myeloid cells are shown. **(B)** CD16a expression levels on total CD16a+, non-classical, and intermediate monocyte populations. **(C)** Mean ROC response and the area under the curve (AUC) with its standard deviation obtained using random forest classifier with 6-fold cross validation in 2 outpatient cohorts using FcγR expression levels on myeloid cells. **(D)** Radar plots summarizing the various features of IgG1-CD16a signaling axis in progressors and non-progressors. Significant differences between the two groups are indicated with asterisks in the radar plot for progressors. P values in (A-B) were calculated using unpaired t-tests. (*P < .05, **P < .01, ***P < .001, ****P < 0.0001).

### mRNA vaccination elicits the production of neutralizing IgG with glycoforms that are distinct from those elicited by infection

We next sought to compare the quality of antibodies produced after SARS-CoV-2 mRNA vaccination and infection. To do so, we studied the antibodies elicited after 1 and 2 doses of the Pfizer BNT162b2 SARS-CoV-2 mRNA vaccine in a group of healthy SARS-CoV-2-naïve adults (Stanford adult vaccine cohort, n=29) (Table S4). Neutralizing titers increased between the post-primary vaccination timepoint (21 days post-dose 1 (PD1)) and the post-boost timepoint (21 days post-dose 2 (PD2)). In all, two doses of mRNA SARS-CoV-2 vaccine elicited robust neutralizing antibody responses that were elevated over peak outpatient neutralizing titers (day 28 shown) (Fig. 3A). Over time after vaccination, the distribution of anti-spike IgG subclasses shifted to a more dominant proportion of IgG1 antibodies (Fig. 3B).

**Figure 3.**
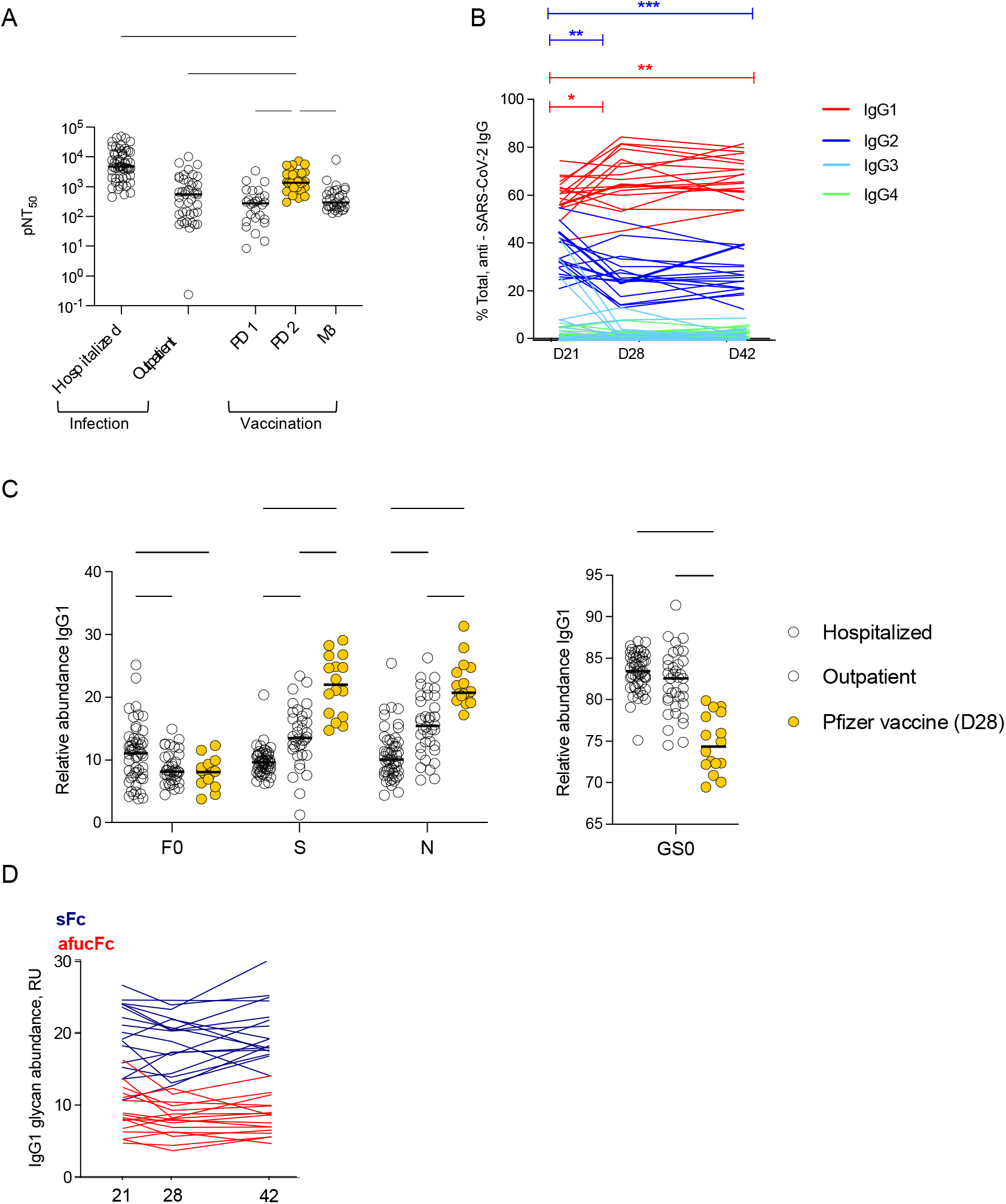
mRNA vaccination elicits high neutralizing antibody titers with Fc glycoforms distinct from both infection-induced phenotypes. **(A)** The half-maximal SARS-CoV-2 pseudovirus neutralizing titers (pNT_50_) in healthy adults following mRNA vaccination (yellow n=29) or in COVID-19 outpatients on study day 28 (blue n=42). PD1: post-dose 1, PD2: post-dose 2, M3: month 3. **(B)** Longitudinal analysis of IgG subclasses on day 21, 28 or 42 post-primary vaccination (n=17). **(C)** SARS-CoV2 IgG1 Fc posttranslational modifications in hospitalized COVID-19 patients (n=52), COVID-19 outpatients (day 28 n=36) and in subjects who received the mRNA SARS-CoV-2 vaccine (day 28 post primary vaccination, n=17). F0: afucosylation, S: sialylation, N: bisection, GS0: galactosylation. The median values have been depicted with a black line. **(D)** Longitudinal analysis of anti-SARS-CoV-2 IgG1 Fc afucosylation (afucFc, red line) and sialylation (sFc, blue line) on day 21, 28 or 42 post-primary vaccination. P values in (A) were calculated using Kruskal Wallis test with Dunn’s correction, (B) using mixed effect analysis with Geissser-Greenhouse and Tukey’s corrections, (C) two-way ANOVA and one-way ANOVA with Tukey’s correction. *P < 0.05; **P < 0.01; ***P < 0.001; ****P < 0.0001.

We next characterized Fc glycoforms of anti-spike IgG to determine whether infection- and mRNA vaccine-elicited IgG were distinct in this respect (*10*). For this analysis, IgG from day 28 of the outpatient COVID-19 study (from non-progressors) were compared to samples drawn from vaccine recipients on day 28 post-primary vaccination (7 days post-boost). Additionally, we compared these groups to IgG from the hospitalized COVID-19 cohort (Mount Sinai). Levels of IgG1 Fc afucosylation were similar between the outpatient and vaccine-elicited IgG (Fig. 3C) and both groups were significantly reduced in afucosylation relative to hospitalized patients. Interestingly, vaccination-induced IgG was significantly enriched in Fc sialylation over both outpatient and hospitalized COVID-19, suggesting differential regulation of Fc sialylation by mRNA vaccination and infection, though we cannot exclude a contribution from demographic features that were not matched between cohorts. The relative homogeneity of Fc glycosylation in response to mRNA vaccination contrasted with the heterogeneity observed in natural infection, as well as with our previous observations after seasonal influenza virus vaccination, suggesting differences in the response that may be based on the context of antigen encounter, antigen experience, or different vaccine platforms (*29*). Vaccine-elicited Fc afucosylation and sialylation were relatively stable over time, similar to the stability observed after natural infection (Fig. 3D, Fig. S1C). Thus, SARS-CoV-2 infection and mRNA vaccination both elicited high neutralizing titers, but distinct and stable levels of IgG1 Fc afucosylation and sialylation.

### Afucosylated immune complexes trigger inflammation in the lung *in vivo*

To study the functional relevance of the differential glycosylation of mRNA- and infection-elicited IgG, we established an *in vivo* experimental model designed specifically to enable dissection of human antibody signaling outcomes in the lung, in the absence of any additional effects imposed by infection. In this model, pre-formed human IgG ICs, simulating what would be formed during an infection, are delivered to lungs of mice that express human, instead of murine, FcγRs, with cell-specific distribution that mimics the human system (*14*). Polyclonal IgG pools were generated from purified serum IgG. Pools were from patients with elevated (pool 1, >20%) or normal levels (pool 2, <10%) of afucosylated IgG or from sera of mRNA-vaccinated adults (pool 3). Pools 1 and 2 did not significantly differ in other glycan modifications, and all 3 pools exhibited comparable distribution of IgG subclasses, though we acknowledge that pool 3 trends toward a lower IgG1 and higher IgG2 content (Fig. S4A, S4B). All IgG pools were standardized for binding to SARS-CoV-2 spike (Fig. S4C). Mice were intratracheally administered ICs composed of the anti-SARS-CoV-2 IgG and trimeric SARS-CoV-2 spike protein. Four hours following IC administration, contents of BAL fluid were analyzed for immune cells and soluble factors. This system provided a context in which to specifically study how modulation of IgG Fc-FcγR interactions impacts the immune response in the lung.

BAL fluid collected from the lungs of mice that were treated with afucosylated ICs (pool 1) had significantly elevated frequencies of neutrophils and monocytes over BAL fluid from mice treated with fucosylated ICs (pool 2) or mRNA-vaccine elicited ICs (pool 3) (Fig. 4A, Fig. S5). BAL fluid from mice that received afucosylated ICs was also distinguished from all other experimental conditions by increased concentrations of proinflammatory cytokines and chemokines. TNFα, IL-6, CCL3, CCL4, CXCL1, and CXCL10 were significantly and uniquely upregulated while no difference was observed in levels of the immunoregulatory or immunosuppressive cytokine IL-10 (Fig. 4B). Collectively, these findings functionally distinguish afucosylated IgG, characteristic of severe COVID-19, from the highly sialylated and fucosylated, vaccine-elicited antibody glycoforms *in vivo*.

**Figure 4.**
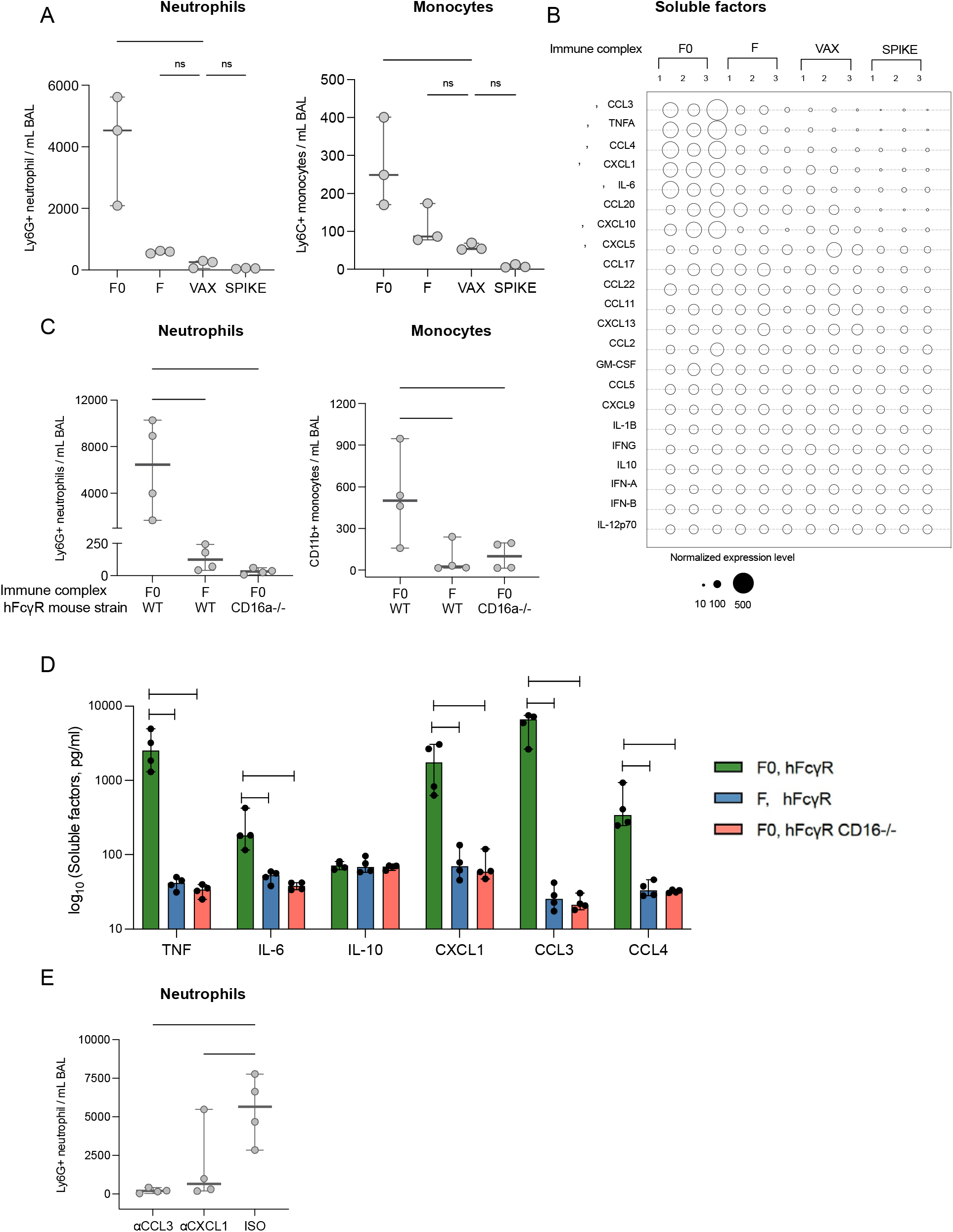
Afucosylated IgG immune complexes promote cell infiltration and proinflammatory cytokine production *in vivo*. **(A)** Immune cells measured in the bronchoalveolar lavage (BAL) fluid of human Fcγ receptor mice (HFcγR) that were treated with either afucosylated (F0, pool 1), normally fucosylated (F, pool 2), vaccine-induced (VAX, pool 3) immune complexes or spike alone, by intratracheal administration. **(B)** Cytokine/chemokine levels in the BAL of the various groups of mice. The size of the bubble represents normalized cytokine and chemokine levels. p values are indicated for each soluble factor (blue: F0 vs F, red: F0 vs VAX). **(C)** Immune cell subsets, **(D)** cytokines (TNF, IL-6, IL-10) and chemokines (CXCL1, CCL3, CCL4) quantitated in the bronchoalveolar lavage (BAL) fluid of hFcγR or hFcγR mice with a specific deletion in CD16a (CD16a-/-) that received afucosylated (F0, pool 1) or normally fucosylated (F, pool 2) immune complexes by intratracheal administration. In **(A)** and **(C)** neutrophils were defined as Ly6G+ CD11b+ CD2- CD3- B220- cells and total monocytes defined as CD11b+ Ly6G- MERTK- MHC IA/IE- CD2- CD3- B220- cells. **(E)** Frequency of Ly6G+ CD11b+ CD2- CD3- B220- neutrophils in BAL fluid of hFcγR mice that were pre-treated with chemokine neutralizing mAbs (αCXCL1 and αCCL3) or isotype control followed by administration of afucosylated immune complexes. The median and the 95% confidence interval have been depicted in each graph with solid lines. P values in (A-D) were calculated using one-way ANOVA with Dunnett’s correction using n=3 mice per group for A and B and n=4 mice per group for C-E. Data in A-E are representative of at least two independent experiments. *P < 0.05; **P < 0.01; ***P < 0.001; ****P < 0.0001.

We next assessed the FcγR dependence of the immune response to afucosylated ICs. Mice specifically lacking expression of CD16a (CD16a-/-), but expressing all other human FcγRs, did not exhibit a similar inflammatory response to afucosylated ICs as mice expressing the complete human repertoire (WT) (Fig. 4C, D). This showed that the inflammatory potential of afucosylated IgG1 was almost entirely dependent on the presence of CD16a-expressing immune effector cells. Because CXCL1 and CCL3 are known neutrophil chemoattractants, we next asked whether these molecules mediated the neutrophil influx after afucosylated IC administration. Indeed, pre-administration of blocking monoclonal antibodies against the chemokines CXCL1 or CCL3 led to a significant reduction in neutrophil recruitment (Fig. 4E) (*30*, *31*). Together, these findings support a mechanism in which afucosylated ICs in the lung trigger CD16a-dependent production of chemokines which promote subsequent influx of innate immune cells.

## Discussion

Prognostic biomarkers and treatments that may halt the progression to severe COVID-19 are urgently needed to prevent mortality associated with this disease. To identify new avenues of treatment, mechanisms underlying the distinct trajectories in COVID-19 must be clarified. Here, we show that early antibody quality and the expression of cognate FcγRs on peripheral monocytes may be used to anticipate distinct COVID-19 trajectories, including progression to more severe outcomes. Overall, mild COVID-19 patients who experienced a worsening disease trajectory were characterized by the absence of an early robust neutralizing antibody response with elevations in both afucosylated anti-spike IgG and the CD16a receptor on myeloid cells. The IgG elicited by SARS-CoV-2 infection was heterogenous in Fc glycosylation relative to IgG generated in response to SARS-CoV-2 mRNA vaccination. Vaccine-elicited IgG exhibited high neutralization and low afucosylation, along with other significant differences in Fc glycoforms. The early inflammatory response to ICs in the lung was a function of the abundance of IgG1 afucosylation and CD16 expression.

Our data support a model in which the combination of a lack of early SARS-CoV-2 neutralization and an enhanced afucosylated IgG-CD16a signaling axis contribute to the inflammatory phenotype of severe COVID-19. We propose that this may be one mechanism contributing to the hyperinflammatory response in severe COVID-19. Determining how various immune aberrancies, including those described here and others such as elevated IL-6 or an impaired renin-angiotensin system might contribute to the pathogenesis of severe COVID-19 will require the development of new animal models (*2, 24, 32, 33*). To examine whether afucosylated ICs can augment the inflammatory milieu in the lungs, we established a model to specifically evaluate the impact of human IgG signaling on the pulmonary inflammatory response. This model advances our ability to evaluate human IgG antibodies in a functional dimension, beyond what *in vitro* approaches can reveal. We show that the afucosylated IgG/CD16a signaling axis can result in a complete remodeling of the inflammatory lung milieu. Of note, the increased frequency of neutrophils and monocytes observed within the lungs of mice that received afucosylated ICs mirrors what has been observed in some severe COVID-19 patients (*4, 21, 34, 35*). Tissue-resident alveolar macrophages likely serve as an initial effector of afucosylated IC activity in this model as they are the predominant innate immune cell population within the lung, express high levels of CD16a, and can produce many of the observed soluble factors (*11*). This *in vivo* model is not a model of COVID-19 pathogenesis; rather, it enables a more targeted investigation of how distinct human antibody repertoires activate effector cells and the complex molecular changes involved in those interactions specifically within the lung. Animal models that more accurately reflect the immunophenotype of patients at highest risk for mortality in COVID-19 are needed to truly study the pathogenesis of this disease.

While we did not observe a significant correlation between IgG afucosylation and the demographic features studied here, it is known that IgG post-translational modifications are associated with specific patient characteristics including sex and age (*36*). Thus, differences in demographics between our cohorts may have contributed to our findings. How IgG glycosylation is regulated is not fully understood, but numerous studies support a role for both heritable and non-heritable influences (*29, 37–41*). We believe that the data in our current manuscript are the first to support a direct role for plasmablast FUT8 expression as a determinant of IgG afucosylation. Defining specific regulatory pathways of IgG glycosylation will be important for modulating the *in vivo* activities of IgG to improve human disease outcomes.

## Materials and Methods

### Clinical cohorts and samples

Characterization of these samples at Stanford was performed under a protocol approved by the Institutional Review Board of Stanford University (protocol #55718).

### Stanford Lambda cohort (Cohort 1)

120 participants were enrolled in a phase 2 randomized controlled trial of Peginterferon Lambda-1a beginning April 25, 2020 (Lambda, NCT04331899). Inclusion/exclusion criteria and the study protocol for the trial have been published(*15*). Briefly, adults aged 18-75 years with an FDA emergency use authorized reverse transcription-polymerase chain reaction (RT-PCR) positive for SARS-CoV-2 within 72 hours prior to enrollment were eligible for study participation. Exclusion criteria included hospitalization, respiratory rate >20 breaths per minute, room air oxygen saturation <94%, pregnancy or breastfeeding, decompensated liver disease, recent use of investigational and/or immunomodulatory agents for treatment of COVID-19, and prespecified lab abnormalities. All participants gave written informed consent, and all study procedures were approved by the Institutional Review Board of Stanford University (IRB-55619). Participants were randomized to receive a single subcutaneous injection of Lambda or saline placebo. Peripheral blood was collected at enrollment, day 5, and day 28 post enrollment. A subset of participants (n=80) returned for long-term follow-up visits 4-, 7-, and 10-months post enrollment, with peripheral blood obtained. Longitudinal samples from the 56 SARS-CoV-2-infected outpatients who were in the placebo arm of the broader Lambda study were obtained and assessed here.

### Stanford Favipiravir Cohort (Cohort 2)

149 participants were enrolled in a phase 2 randomized controlled trial of Favipiravir beginning July 12, 2020 (NCT04346628). Inclusion/exclusion criteria and the study protocol for the trial are publicly available. Briefly, adults aged 18 – 80 years old were enrolled within 72 hours of a positive NAAT for SARS-CoV-2. Upon enrollment, participants were mildly symptomatic with no evidence of respiratory distress. Participants were randomized to receive favipiravir or placebo. Participants were followed for 28 days, with study visits on days 1, 5, 10, 14, 21 and 28. At each study visit, clinical assessment was performed and oropharyngeal swabs and blood samples were collected. Samples collected upon enrollment from the 69 SARS-CoV-2-infected outpatients who were in the placebo arm of the broader Favipiravir study were obtained and assessed here.

### Mount Sinai Cohort (Hospitalized COVID-19 patients)

Fifty-two samples were obtained from hospitalized COVID-19 patients enrolled in the Mount Sinai Health System (MSHS) collected by the Mount Sinai COVID-19 biobank ^22^. The median age was 65 years old with a range from 33 - 98 years old. There are 31 males and 21 females in the study. 13 patients succumbed to disease.

### Stanford Adult vaccine cohort

Fifty-seven healthy volunteers were enrolled in the study approved by Stanford University Institutional Review Board (IRB 8629). The median age was 36 years old with a range from 19 - 79 years old. There are 28 males and 29 females in the study. There are 27 White participants, 23 Asian participants, 4 Black participants, 1 Native American participant, and 2 other participants.

#### Cell lines

Human embryonic kidney (HEK) 293T (American Type Culture Collection, ATCC; CRL-3216) and Vero (ATCC; CCL-81) cells were grown and maintained in 1X Dulbecco’s Modified Eagle Medium (DMEM; ThermoFisher Scientific) supplemented with 10% fetal bovine serum (FBS).

#### Cloning, expression, and protein purification

The His_6_-tagged SARS-CoV-2 RBD and full-length SARS-CoV-2 spike protein were purified in house as previously described.^3^ Briefly, both the constructs were transiently transfected into Expi293F cells (Thermo Fisher Scientific) and proteins purified from culture supernatants Ni-nitriloacetic acid (NTA) resin (GE HealthCare).

#### Generation of SARS-CoV-2 pseudoparticles

To generate vesicular stomatitis virus (VSV) pseudotyped with the S of SARS-CoV-2, we first constructed an expression plasmid encoding the Wuhan S. We did this by modifying a pCAGGS mammalian expression vector encoding the full-length Wuhan S and deleting its last 18 amino acids of the cytoplasmic domain, which we call pCAGGS-SΔ18. This reagent was produced under HHSN272201400008C and obtained through BEI Resources, NIAID, NIH: Vector pCAGGS containing the SARS-related coronavirus 2, Wuhan S, NR52310. To generate VSV pseudotyped with SARS-CoV-2 S, we first coated 6-well plates with 0.5 μg/mL poly-D-lysine (ThermoFisher, Cat. No. A3890401) for 1 to 2 hours at room temperature (RT). After poly-D-lysine treatment, plates were washed three times with sterile water and then seeded with 1.5e6 cells of HEK 293T per well. After 24 hours (h), cells were transfected with 1 μg of pCAGGS-SΔ18 per well using Lipofectamine 2000 transfection reagent (ThermoFisher, Cat. No., 11668019). 48 h after transfection, the cells were washed once with 1X phosphate buffered saline (PBS), and were infected with VSV-ΔG-GFP/nanoluciferase (a generous gift from Matthias J. Schnell) at a multiplicity of infection of 2 to 3 in a 300 μL volume. Cells were infected for an hour with intermittent rocking every 15 minutes. After infection, the inoculum was carefully removed, and the cell monolayer was washed three times with 1X PBS to remove residual VSV-ΔG-GFP/nanoluciferase. Two mL of infection media (2% FBS, 1% glutamine, 1% sodium pyruvate in 1X DMEM) was added to each well. At 24 h post-infection, the supernatants from all the wells were combined, centrifuged (600 *g* for 10 min, 4°C), and stored at −80°C until use.

#### Neutralization assays

Vero cells were seeded at 5e5 cells per well in 50 μL aliquots in half area Greiner 96-well plates (Greiner Bio-One; Cat. No. 675090) 24 h prior to performing the neutralization assay. On separate U-bottom plates, patient plasma was plated in duplicates and serially 5-fold diluted in infection media (2% FBS, 1% glutamine, 1% sodium pyruvate in 1X DMEM) for a final volume of 28 μL per well. We also included ‘virus only’ and ‘media only’ controls. Twenty-five microliters of SARS-CoV-2 pseudo-typed VSV particles (containing 500 to 1500 fluorescent forming units) were added to the wells on the dilution plate, not including the “virus-free” column of wells and incubated at 37°C for 1 hour. Prior to infection, Vero cells were washed twice with 1X PBS and then 50 μL of the incubated pseudo-typed particles and patient plasma mixture was then transferred from the U-bottom 96-well dilution plates onto the Vero cells and placed into an incubator at 37°C and 5% CO2. At 24 h post-incubation, the number of GFP-expressing cells indicating viral infection were quantified using a Celigo Image Cytometer. We first calculate percent infection based on our ‘virus only’ controls and then calculate percent inhibition by subtracting the percent infection from 100. A non-linear curve and the half-maximal neutralization titer (pNT_50_) were generated using GraphPad Prism.

#### ELISA

ELISA was performed following a modified version of a protocol described previously3. Briefly, 96 Well Half-Area microplates (Corning (Millipore Sigma)) plates were coated with antigens at 2μg/ml in PBS for 1h at room temperature (RT). Next, the plates were blocked for an hour with 3% non-fat milk in PBS with 0.1% Tween 20 (PBST). All serum samples from patients with COVID-19, and the negative controls were heated at 56°C for 1h, aliquoted and stored at −80°C. Sera were diluted fivefold starting at 1:50 in 1% non-fat milk in PBST. 25μl of the diluted sera was added to each well and incubated for 2h at RT. Following primary incubation with the sera, 25 μl of 1:5000 diluted horse radish peroxidase (HRP) conjugated anti-Human IgG secondary antibody (Southern Biotech) was added and incubated for 1h at RT. The plates were developed by adding 25μl/well of the chromogenic substrate 3,3’,5,5’-tetramethylbenzidine (TMB) solution (Millipore Sigma). The reaction was stopped with 0.2N sulphuric acid (Sigma) and absorbance was measured at 450nm (SPECTRAmax iD3, Molecular Devices). The plates were washed 5 times with PBST between each step and an additional wash with PBS was done before developing the plates. All data were normalized between the same positive and negative controls and the binding AUC has been reported.

#### IgG Fc glycan analysis

Methods for relative quantification of Fc glycans and IgG subclasses have been previously described^20,23^. Briefly, IgG were isolated from serum by protein G purification. Antigen-specific IgG were isolated on NHS agarose resin (ThermoFisher; 26196) coupled to the protein of interest. Following tryptic digestion of purified IgG bound to antigen-coated beads, nanoLC-MS/MS analysis for characterization of glycosylation sites was performed on an UltiMate3000 nanoLC (Dionex) coupled with a hybrid triple quadrupole linear ion trap mass spectrometer, the 4000 Q Trap (SCIEX). MS data acquisition was performed using Analyst 1.6.1 software (SCIEX) for precursor ion scan triggered information dependent acquisition (IDA) analysis for initial discovery-based identification.

For quantitative analysis of the glycoforms at the N297 site of IgG1, multiple-reaction monitoring (MRM) analysis for selected target glycopeptides and their glycoforms was applied using the nanoLC-4000 Q Trap platform to the samples which had been digested with trypsin. The m/z of 4-charged ions for all different glycoforms as Q1 and the fragment ion at m/z 366.1 as Q3 for each of transition pairs were used for MRM assays. A native IgG tryptic peptide (131-GTLVTVSSASTK-142) with Q1/Q3 transition pair of, 575.9^+2^/780.4 was used as a reference peptide for normalization. IgG subclass distribution was quantitatively determined by nanoLC-MRM analysis of tryptic peptides following removal of glycans from purified IgG with PNGase F. Here the m/z value of fragment ions for monitoring transition pairs was always larger than that of their precursor ions with multiple charges to enhance the selectivity for unmodified targeted peptides and the reference peptide. All raw MRM data was processed using MultiQuant 2.1.1 (SCIEX). All MRM peak areas were automatically integrated and inspected manually. In the case where the automatic peak integration by MultiQuant failed, manual integration was performed using the MultiQuant software.

#### Immune cell phenotyping and FcγR quantification

Cryopreserved human PBMCs collected upon enrollment on study day 0 were rapidly thawed, washed, and blocked with Human TruStain FcX (BioLegend) to reduce nonspecific binding. Cells were then stained for viability with Live/Dead Fixable Staining Kit (ThermoFisher) as well as the following antibodies: anti-CD3 (clone OKT3), anti-CD11c (clone S-HCL-3), anti-CD14 (clone M5E2), anti-CD16 (clone 3G8), anti-CD19 (clone SJ25C1), anti-CD21 (Bu32), anti-CD27 (clone O323), anti-CD32 (STEMCELL Technologies; clone IV.3), anti-CD32B/C (clone S18005H), anti-CD38 (clone S17015A), anti-CD56 (clone 5.1H11), anti-CD138 (clone IA6-2), anti-FucT-VIII (Santa Cruz Biotechnologies; clone B-10), anti-HLA-DR (clone L243) purchased from BioLegend unless noted otherwise. After staining, cells were fixed and acquired using an Attune NxT flow cytometer (Invitrogen). In the case of intracellular anti-FucT-VIII staining, cells were further permeabilized using Intracellular Staining Permeabilization Wash Buffer (BioLegend) prior to acquisition by flow cytometry. Bulk myeloid cells were defined as viable CD3- CD19- CD56- CD11c+ HLA-DR+ cells, while CD16a+ monocytes within this population were additionally positive for CD16a (Fig S3). Within CD16a+ monocytes, non-classical (NC) monocytes were CD16a+ CD14- while intermediate (int) monocytes were CD16a+ CD14+. Leukocyte expression of FcγRs was quantified by measuring the median fluorescence intensity (MFI) of a particular FcγR and comparing it to the MFI of stained Quantum™ Simply Cellular microsphere beads (Bangs Laboratories) of known and discrete antibody-binding capacities. Total CD19+ B cells were similarly assessed from within viable PBMCs. Plasmablasts were further defined as CD19+ CD27+ CD38++. Memory B cells were defined as CD19+ CD27+ IgD-, double negative (DN) B cells were CD19+ CD27- IgD-, and naïve B cells were CD19+ CD27- IgD+.

#### *In vivo* lung inflammation model

All *in vivo* experiments were performed in compliance with federal laws and institutional guidelines and have been approved by the Stanford University Institutional Animal Care and Use Committee. Polyclonal IgG was isolated from PCR-positive SARS-CoV-2 patients’ plasma samples, pooled based on the frequency of afucosylated anti-RBD IgG1 (>20% or <10%). Similarly, plasma from all vaccinated patient samples were pooled and IgG was purified. The purified IgG pools were incubated with SARS-CoV-2 spike trimer at a 20:1 molar ratio overnight at 4°C. Immune complexes were intratracheally administered to 8–12-week-old humanized FcγR or CD16a-deficient mice. Experimental groups were consistently matched for sex and age. 4hr post-administration, mice were sacrificed and bronchoalveolar lavage (BAL) was performed. Immune cells were isolated from within the BAL fluid and stained with the following cell staining panel: Live/Dead Fixable Dye (ThermoFisher), anti-CD2 (BioLegend; clone RM2-5), anti-CD3 (BioLegend; clone 17A2), CD11b (BioLegend; clone M1/70), anti-CD45 (BioLegend; clone I3/2.3), anti-human CD64 (BioLegend; clone 10.1), anti-B220 (BioLegend), anti-anti-Ly6G (BioLegend; clone 1A8), anti-MERTK (BioLegend; clone 2B10C42), anti-MHC II (BioLegend; clone M5/114.15.2). Once stained, cells were fixed and acquired via an Attune NxT flow cytometer (Invitrogen). Neutrophils were defined as viable Ly6G+ CD11b+ CD3- B220- leukocytes. Monocytes were defined as viable CD11b+ Ly6G- MERTK- MHC IA/IE- CD3- B220- leukocytes (Fig S6). Cell-free BAL fluid was stored at 4°C and processed within 24hr to measure cytokine/chemokine content using a LEGENDplex bead array kit (BioLegend).

#### *In vivo* blocking experiments

In chemokine-blockade experiments, mice received intraperitoneal injections of 5mg/kg anti-CXCL1, anti-CCL3, or rat IgG2a isotype control (MAB453, MAB4502, MAB006; R&D) 8hr prior to immune complex administration and immune complex administration and BAL were performed as described above.

#### Data and Statistical analysis

The log10+1 transformed half-maximal serum neutralization titers (pNT_50_) were used to generate the heatmap.

Python version 3.8.5 was used for machine learning. The class progressor was mapped to 1 and non-progressor was mapped to 0, making it a binary classification problem.

To determine whether the combination of low/no neutralizing antibodies and elevated IgG Fc afucosylation was a predictor of worsening disease trajectory, a logistic regression model was used. The model was trained using data from Cohort 1 (training set), and to obtain the best hyperparameters, GridSearch cross-validation (cv) was performed. The model was tested using an independent test set (Cohort 2) and the ROC AUC score was generated.

To generate ROC AUC scores from FcγRs frequency and expression to distinguish progressors and non-progressors, Random Forest Classifier was used. The input data was split using 6-fold cross validation in which the classifier was trained on 5 folds of the data and tested on the remaining part of the data. The ROC response for all these different datasets were used for calculating the mean area under curve.

R Studio (version 1.2.1335) was used to perform the multivariate regression analyses and to generate the radar plots and bubble plot using ggplot2 package. For the radar plots, each feature was normalized across the entire dataset and the mean value within each cohort (progressor and non-progressor) was plotted. For the bubble plot, cytokine and chemokine levels were normalized between 0 and the average of all values across all the groups.

All other data were analyzed with GraphPad Prism 9.0 software.

## Acknowledgments

We thank Stanford CTRU Biobank, Catherine A. Blish, Hector Bonilla, Karen Jacobson, Diego Martinez Mori, Kattria van der Ploeg, Sharon Chinthrajah, Tina Sindher, Will Collins, James Liu, Joe G, Anthony Buzzanco, Katia Tkachenko, Mihir Shah, Allie Lee, Kathleen Jia, Eric Smith, Iris Chang, Evan Do and Diane Dunham for support with clinical protocol, patient care, and/or collection/provision of patient samples. Financial support from Stanford’s Innovative Medicines Accelerator and operational support from Stanford ChEM-H is acknowledged.

## Author contributions

SC, JG, GST, TTW designed the study and analyzed data. SC, JG, GST, TTW wrote the manuscript with input from EMD, KCN, TUM, PJ, VM, and UA. SC, JG, BLS, GST, VM, SC, MD, UA, AS, BY-LC, KQTT, CS, RS, SZ and XJ performed experiments. AC, STC, TG, FG, YG, SK, MP, SLG, SG, TUM, MM, SDB, MMD, MH, CK, HM, YM, EDM, KCN, BP, US, AS, and PJ provided critical support and/or reagents.

## Supplemental information

Correspondence and requests for materials should be addressed to T.T.W

## Data availability

All raw data are available in the manuscript or from the corresponding author on request.

## Supplemental Figures

**Supplemental Figure S1:**
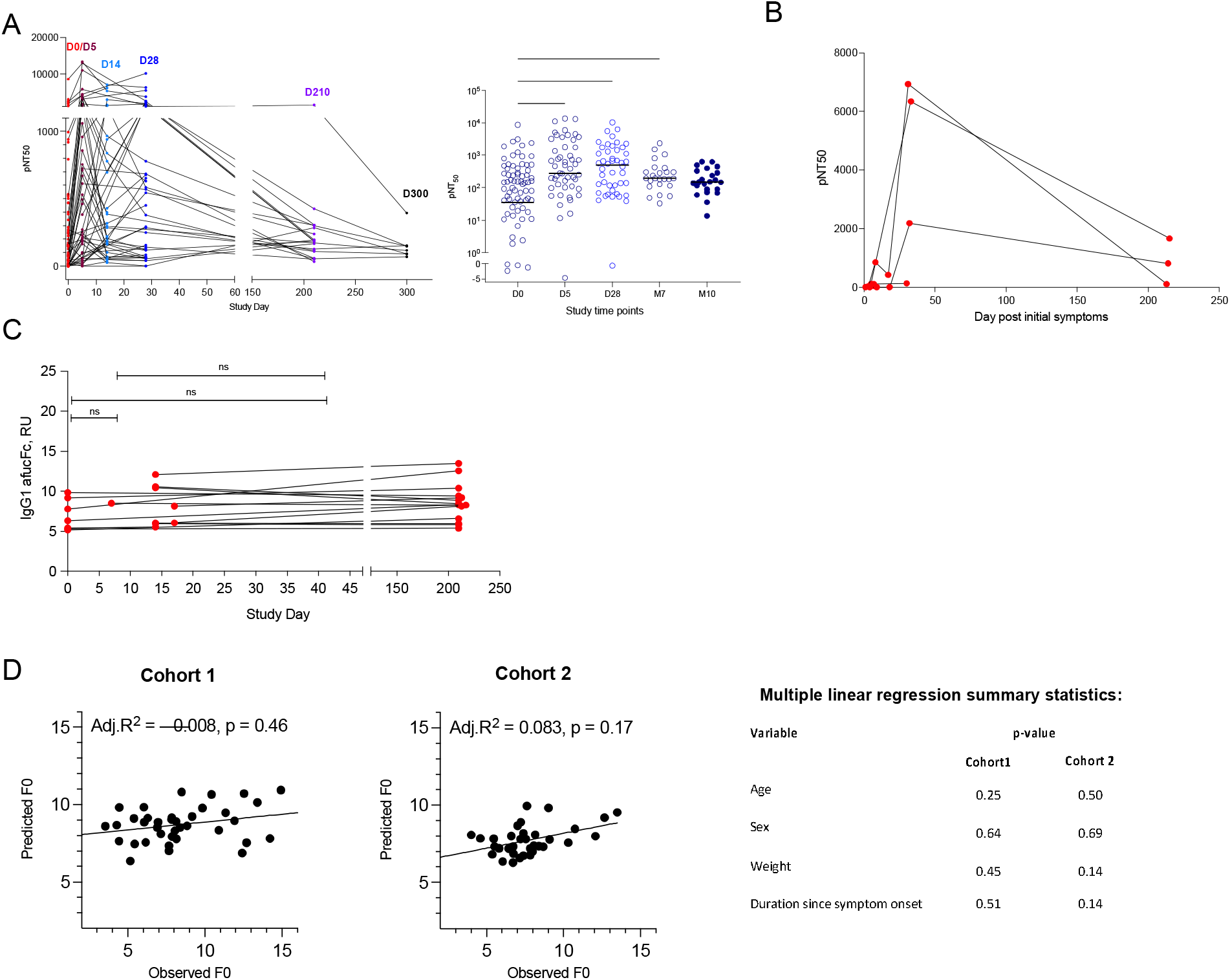
**(A)** Half-maximal SARS-CoV-2 pseudovirus neutralizing titers (pNT_50_) at each study time point. Left panel: graphed to match format of Figure 1A. Right panel: plotted to clearly show the median pNT_50_ at each study time point. **(B)** The kinetics of neutralizing antibody response over time of progressors in Cohort 1. **(C)** Abundance of SARS-CoV-2 specific afucosylated IgG1 in Cohort 1 COVID-19 outpatients over time (n=19). **(D)** Multivariate regression analysis to test the contribution of age, sex, weight, or duration since COVID-19 onset on abundance of afucosylated anti-RBD IgG1 in two cohorts. *P < 0.05; **P < 0.01; ***P < 0.001; ****P < 0.0001

**Supplemental Figure S2:**
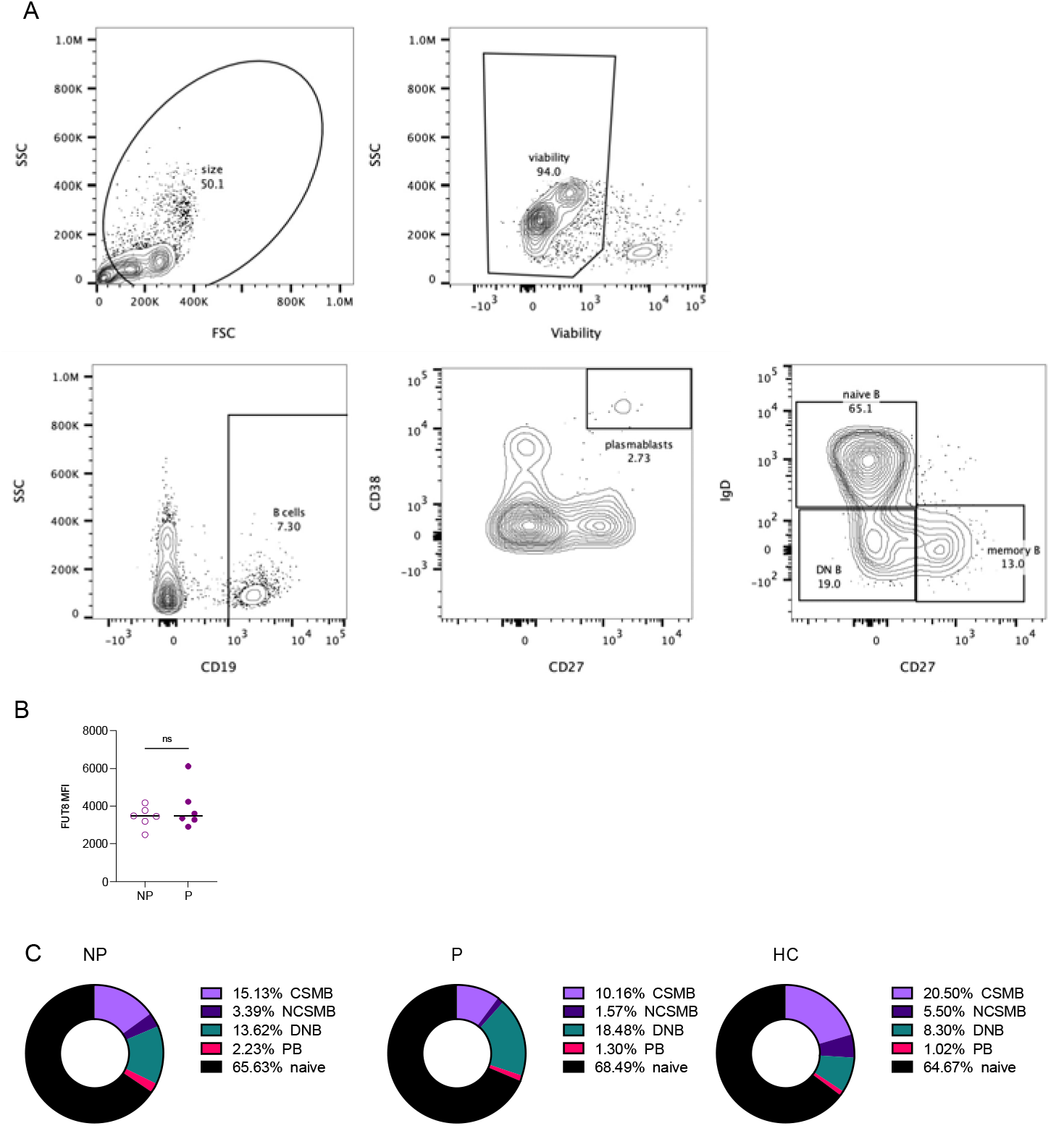
**(A)** Gating strategy for B cell subsets. Total CD19+ B cells were assessed from within viable PBMCs. Plasmablasts were further defined as CD19+ CD27+ CD38++. Memory B cells were defined as CD19+ CD27+ IgD-, double negative (DN) B cells were CD19+ CD27- IgD-, and naïve B cells were CD19+ CD27- IgD+. **(B)** FUT8 expression within total viable PBMCs in progressors (P; n =6) and sex-matched non-progressors (NP; n=6). **(C)** Distribution of B cell subsets (class switched memory B cells (CSMB), nonclass switched memory B cells (NCSMB), CD27- IgD- double negative (DNB), plasmablast (PB), and naïve B cells in progressors (P; n = 6), non-progressors (NP, n = 6), and healthy controls (HC, n = 4). P value in (B) was calculated using unpaired Student’s t test.

**Supplemental Figure S3:**
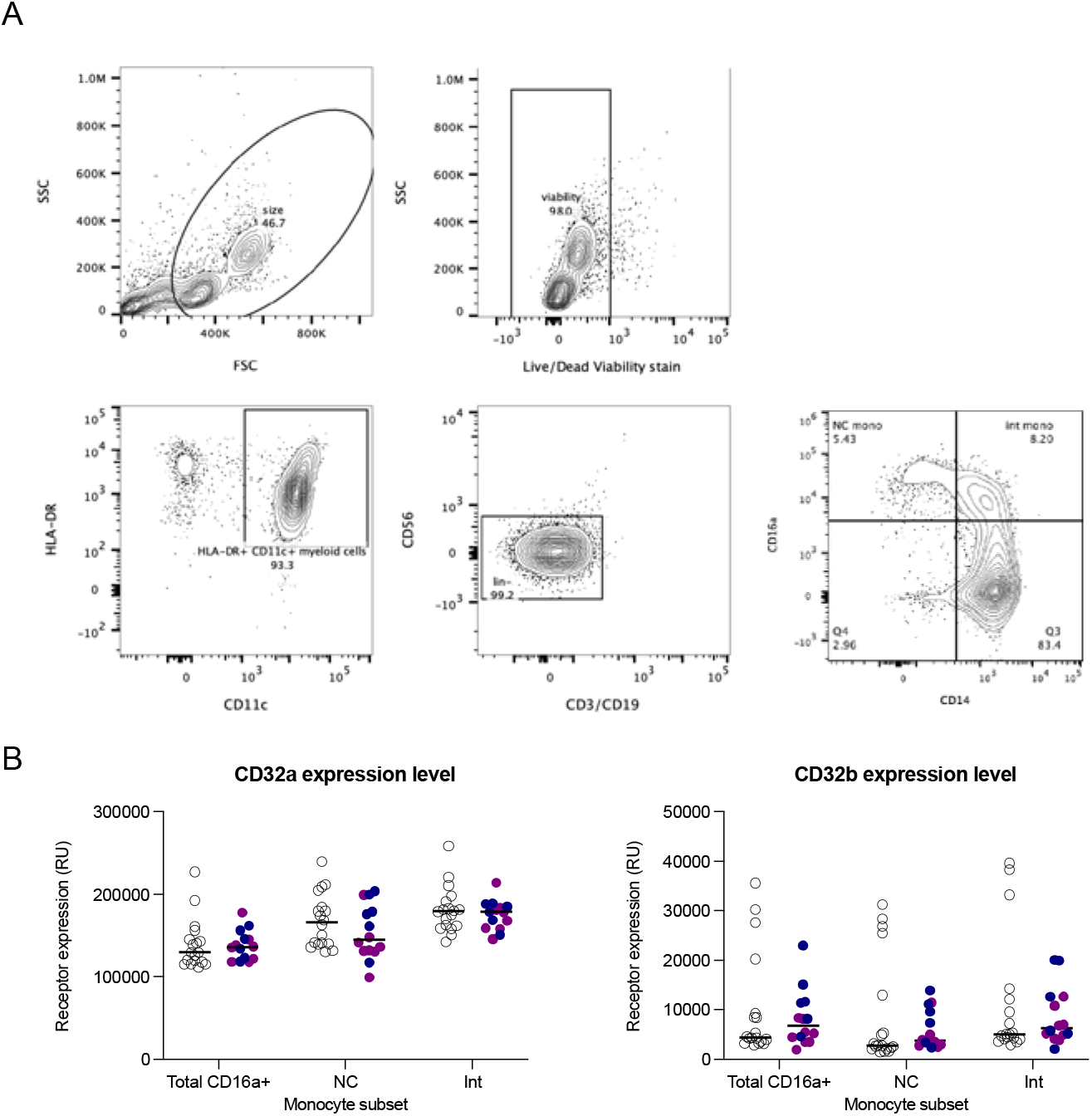
**(A)** Gating strategy for day 0 monocyte subsets and FcγR expression. Bulk myeloid cells were defined as viable CD3- CD19- CD56- CD11c+ HLA-DR+ cells, while CD16a+ monocytes within this population were additionally positive for CD16a. Within CD16a+ monocytes, non-classical (NC) monocytes were CD16a+ CD14- while intermediate (int) monocytes were CD16a+ CD14+. **(B)** FcγR (CD32a and CD32b) expression levels on myeloid cell subsets. P values were determined by unpaired Student’s test.

**Supplemental Figure S4:**
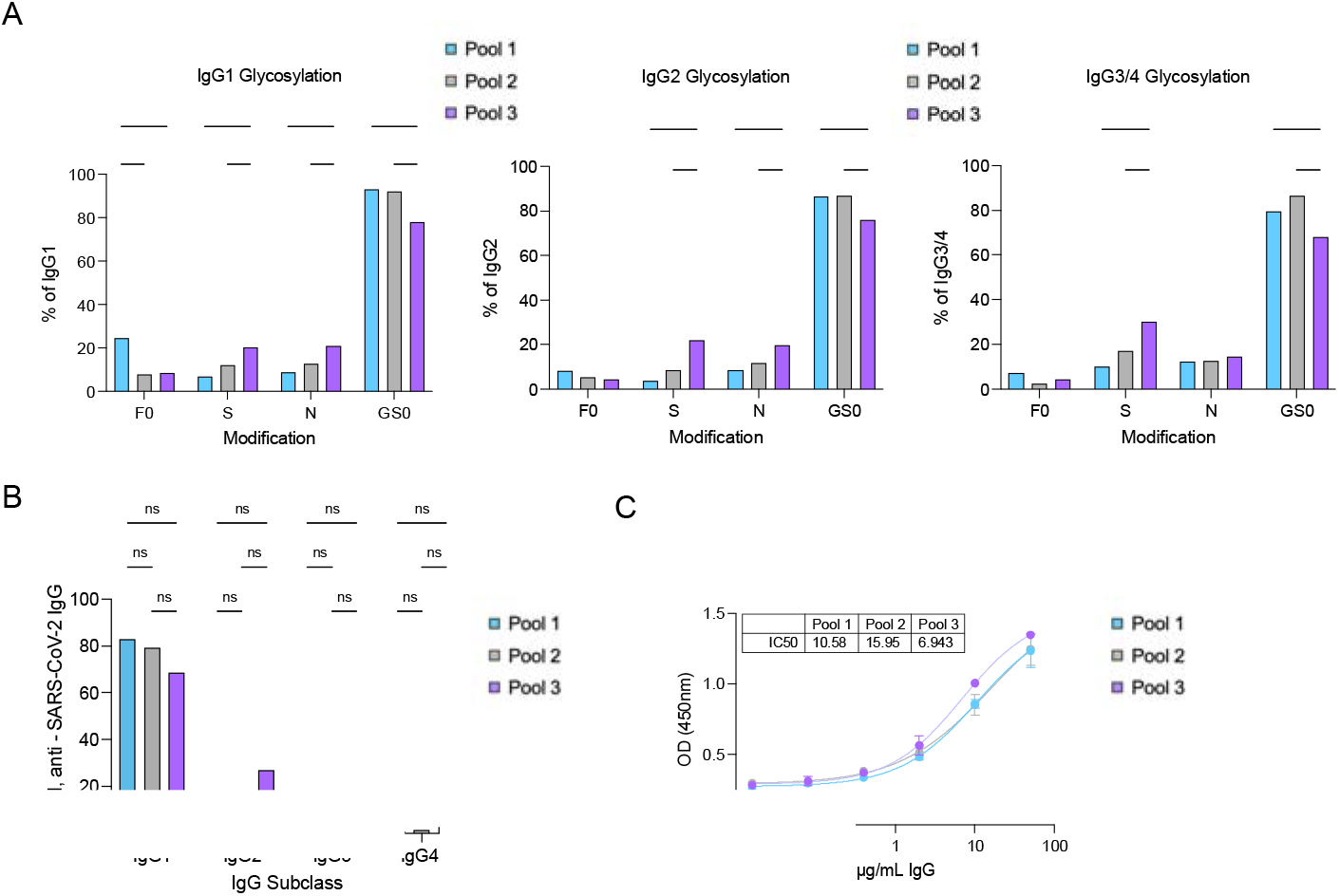
**(A)** Glycan composition (F0-afucosylation, GS0-Galactosylation, S-Sialylation and N-Bisection) of IgG1, IgG2 and IgG3/4 of the individual purified patient IgG comprising polyclonal pool 1 (F0, F0>20%), pool 2 (F, F0<10%), and pool 3 (VAX) polyclonal pools. **(B)** IgG subclass distribution of the individual purified patient IgG comprising polyclonal pool 1, pool 2, and pool 3. **(C)** SARS-CoV-2 full length spike binding of polyclonal pool 1, pool 2, and pool 3. P values in (A-B) were calculated using two-way ANOVA with Šidák’s correction. *P < 0.05; **P < 0.01; ***P < 0.001; ****P < 0.0001

**Supplemental Figure S5:**
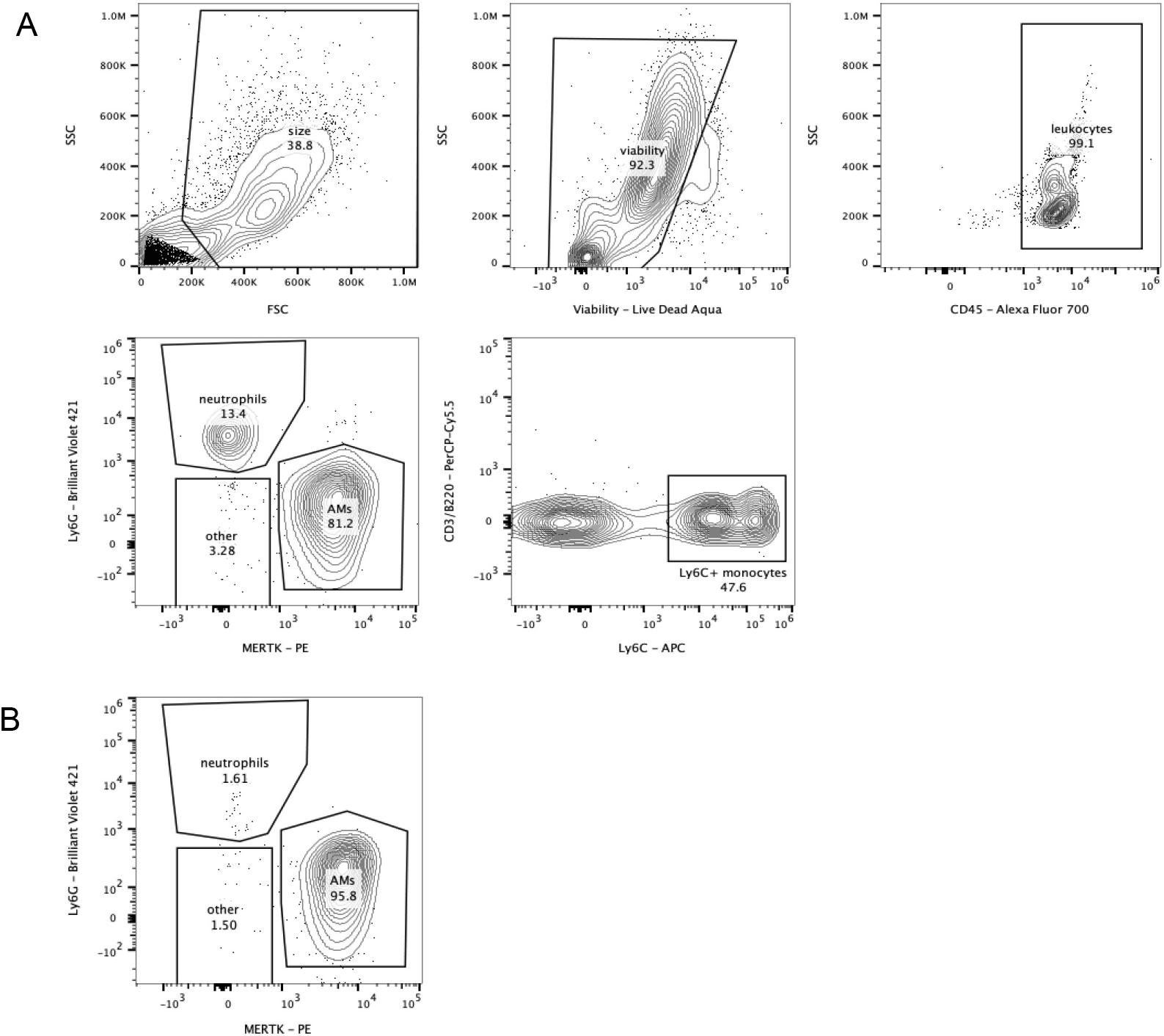
Gating strategy for immune cell infiltrates in mouse BAL. **(A)** Neutrophils were defined as viable Ly6G+ CD11b+ CD3- B220- leukocytes. Monocytes were defined as viable CD11b+ Ly6G- MERTK- MHC IA/IE- CD3- B220- or Ly6C+ CD11b+ Ly6G- MERTK- MHC IA/IE- CD3- B220- leukocytes. **(B)** Alveolar macrophage were magnetically sorted based on positive expression of MERTK. This sorting method consistently resulted in >90% purity as determined by flow cytometry.

## Tables

**Table S1.** Demographics of the COVID-19 infection cohorts.

| Stanford Lambda cohort (Cohort 1) | Non-progressor<br>(n=102) | Missing<br>values | Progressor (n=8) | Missing<br>values | p value |
| --- | --- | --- | --- | --- | --- |
| Age, years (median, 95%CI) | 36 (32.8-38.2) | 0 | 45 (34.4-55.7) | 0 | ns |
| Sex Female | 45 (44.1%) | 0 | 3 (38%) | 0 | ns |
| Race/Ethnicity |  | 0 |  | 0 |  |
| Latinx | 65 (63.7%) |  | 5 (62.5%) |  |  |
| White | 29 (28.4%) |  | 2 (25%) |  |  |
| Asian | 7 (6.9%) |  | 0 (0%) |  |  |
| Others | 1(1%) |  | 1 (12.5%) |  |  |
| Weight (lbs) (median, 95% CI) | 175 (168.1-181.9) | 1 | 198.5 (148.8-248.2) | 0 | ns |
| Body mass index (BMI) (median, 95% CI) | 27.2 (26.5-28.9) | 3 | 28.8 (23.9-33.7) | 0 | ns |
| <25 | 25 (25.3%) |  | 2 (25%) |  |  |
| 25-30 | 38 (38.4%) |  | 3 (37.5%) |  |  |
| >30 | 36 (36.4%) |  | 3 (37.5%) |  |  |
| Median days from symptoms onset to study enrollment | 4 | 17 | 3.5 | 0 | ns |
| Mean days from symptoms onset to study enrollment | 5 | 17 | 3.6 | 0 | ns |
| Median days from randomization to disease resolution | 9 | 0 | 16 | 4 | ns |

| Stanford Favipiravir Cohort (Cohort 2) | Non-progressor<br>(n=67) | Missing<br>values | Progressor (n=7) | Missing<br>values | p value |
| --- | --- | --- | --- | --- | --- |
| Age, years (median, 95%CI) | 40 (36.9-43.1) | 0 | 52 (45.9-58) | 0 | ** |
| Sex Female | 35 (52.2%) | 0 | 3 (42.9%) | 0 | ns |
| Race/Ethnicity |  | 4 |  | 0 |  |
| Latinx | 26 (38.8%) |  | 5 (71.4%) |  |  |
| White | 27 (40.2%) |  | 1 (14.3%) |  |  |
| Asian | 7 (10.4%) |  | 0 (0%) |  |  |
| Others | 3(4.5%) |  | 1 (14.3%) |  |  |
| Weight (lbs) (median, 95% CI) | 180 (170.4-189.6) | 0 | 190 (146.5-233.4) | 0 | ns |
| Body mass index (BMI) (median, 95% CI) | 28.3 (26.8-29.7) | 0 | 30.8 (24.7-37) | 0 | ns |
| <25 | 20 (29.9%) |  | 2 (28.6%) |  |  |
| 25-30 | 21 (31.3%) |  | 1 (14.3%) |  |  |
| >30 | 26(38.8%) |  | 4 (57.1%) |  |  |
| Median days from symptoms onset to study enrollment | 5 | 3 | 5 | 0 | ns |
| Mean days from symptoms onset to study enrollment | 5.7 | 3 | 6.4 | 0 | ns |
| Median days from randomization to disease resolution | 11 | 0 | 16 | 0 | ns |

| Mount Sinai Cohort | Hospitalized<br>(n=52) |
| --- | --- |
| Age, years (median, 95%CI) | 65(60.6-69.4) |
| Sex Female | 21(40.4%) |
| Race/Ethnicity |  |
| Latinx | 13 (25%) |
| White | 7 (13.5%) |
| Asian | 3 (5.8%) |
| Others | 29 (55.8%) |
| Weight (lbs) (median, 95% CI) | N/A |
| Body mass index (BMI) (median, 95% CI) | 26.79(24.6-29.01) |
| <25 | 15(28.8%) |
| 25-30 | 21(40.4%) |
| >30 | 16(30.8%) |
| Median days of sample time point post hospitalization<br>(95% CI) | 4(2.87-5.13) |

P values were calculated by comparing features between the progressors and non-progressors using unpaired t-tests with Welch's correction or Fisher's t-test.

**Table S2.** Description of progressors from Cohort 1.

| Age | Sex | Days of symptoms prior to enrollment | Days to Hospitalization/ED visit post assessment of mild COVID-19 |
| --- | --- | --- | --- |
| 25 | Female | 3 | 0; progressive symptoms leading to hospitalization within 24-hours. |
| 58 | Male | 4 | 5; progressively worsening respiratory symptoms. Referred to ED by study-associated physician on study day 5. |
| 44 | Female | 3 | 1; progressively worsening respiratory symptoms. |
| 34 | Male | 2 | 13; worsening symptoms. |
| 47 | Male | 5 | 3; two ED visits within first 3 days for worsening respiratory symptoms. |
| 59 | Female | 7 | 0; progressive symptoms leading to hospitalization within 24-hours. |
| 28 | Male | 4 | 0; progressive symptoms leading to hospitalization within 24-hours. |
| 46 | Male | 1 | 2; progressive symptoms leading to hospitalization. |

**Table S3.** Description of progressors from Cohort 2.

| Age | Sex | Days of symptoms prior to enrollment | Days to Hospitalization/ED visit post assessment of mild COVID-19 |
| --- | --- | --- | --- |
| 44 | Female | 8 | 8; worsening respiratory symptoms |
| 54 | Male | 2 | 1; sent by study team to ED for worsening respiratory symptoms and leg swelling, resulted in hospitalization |
| 57 | Male | 4 | 2; sent by study team to ED for worsening respiratory symptoms; hospitalized |
| 46 | Male | 5 | 3; worsening symptoms and new rash |
| 63 | Female | 10 | 5; sent by study team to ED for worsening symptoms |
| 52 | Male | 4 | 10; sent by study team to ED for worsening respiratory symptoms; hospitalized |
| 50 | Female | 8 | 3; sent by study team to ED for worsening respiratory symptoms; hospitalized |

**Table S4:** Demographics of the Stanford Adult Vaccination Cohort.

| Characteristics |  |
| --- | --- |
| Age, years (median, 95%CI) | 36(28.3-43.7) |
| Sex Female | 11(64.7%) |
| Race/Ethnicity |  |
| Latinx | 0(0.0%) |
| White | 7(41.2%) |
| Asian | 8(47.1%) |
| Others | 2(11.8%) |

